# Leveraging Large Language Models for Literature-Driven Prioritization of Protein Binding Pockets

**DOI:** 10.1101/2025.05.13.653394

**Authors:** Roman Stratiichuk, Mykola Melnychenko, Ihor Koleiev, Taras Voitsitskyi, Vladyslav Husak, Nazar Shevchuk, Zakhar Osrovsky, Volodymyr Bdzhola, Semen Yesylevskyy, Serhii Starosyla, Alan Nafiiev

**Affiliations:** Receptor.AI Inc., 20-22 Wenlock Road, London N1 7GU, United Kingdom; Department of Biophysics and Medical Informatics, Educational and Scientific Centre “Institute of Biology and Medicine”, Taras Shevchenko Kyiv National University, 64 Volodymyrska Str., 01601, Kyiv, Ukraine; Department of Physics of Biological Systems, Institute of Physics of The National Academy of Sciences of Ukraine, 46 Nauky Ave., 03038, Kyiv, Ukraine; Department of Cellular, Computational and Integrative Biology, The University of Trento, Via Sommarive 9, 38123 Povo (Trento), Italy; Institute of Molecular Biology and Genetics of The National Academy of Sciences of Ukraine, 150 Zabolotnogo Str., 03143, Kyiv, Ukraine; Institute of Organic Chemistry and Biochemistry, Czech Academy of Sciences, CZ-166 10 Prague 6, Czech Republic; Department of Physical Chemistry, Faculty of Science, Palacký University Olomouc, 17. listopadu 12, 771 46 Olomouc, Czech Republic

## Abstract

We present a novel approach for the identification and prioritization of protein binding pockets for small molecules by combining geometric pocket detection with Large Language Models (LLMs). Our method leverages Fpocket to generate candidate pockets, which are then validated against published experimental data extracted from research articles using LLM with a series of prompts fine-tuned to identify and extract residue-level information associated with experimentally confirmed binding sites. We developed a curated benchmark dataset of diverse proteins and associated literature to train and evaluate the LLM’s performance in paper relevance assessment and pocket extraction. The extracted information is then mapped onto protein structures and used to filter and merge the geometry-based predictions, generating a refined volumetric representation of biologically relevant pockets. This hybrid pipeline offers an efficient, accurate and automated method for identifying functional binding pockets, addressing a significant bottleneck in the high-throughput drug discovery workflows. The developed benchmark dataset and methodology are freely available at https://github.com/MelnychenkoM/LLM-benchmark-dataset.

## 1. Introduction

The protein binding pockets are the regions on the protein surface where the ligands — such as small molecules, peptides, nucleic acids, or other proteins — can bind. They play a critical role in facilitating biological processes and are essential for understanding protein function. Accurate identification of these sites is crucial for the development of new drugs and the rational design of novel proteins with specific functions.

Over the years, various computational methods have been developed to predict protein binding pockets. Geometry-based methods rely on the structural identification of cavities or clefts on the protein surface that may accommodate ligands. The most widely used method of this family is Fpocket [1] which uses alpha spheres [2] to represent the shape of the pocket cavities.

Energy-based methods, such as EASYMIFs and SITEHOUND [3], use clustering and filtering of the molecular interaction fields (MIFs) to locate the spatial regions on the protein surface that are energetically favorable for ligand binding. Despite their advantages in accounting for biologically relevant interactions, these techniques are computationally more expensive and require careful parameter tuning.

Methods like ConCavity [4] combine a geometry-based approach with a sequence conservation analysis to prioritize the tentative pockets based on their evolutionary significance. However, such techniques often overlook unique non-conserved binding pockets and perform poorly for the proteins without a large number of homologs.

In recent years pocket detection techniques based on machine learning and deep learning emerged. Such methods as DeepSite [5], DeepPocket [6], PUResNet [7] and PUResNetV2.0 [8] integrate diverse geometric, energetic and evolutionary features resulting in improved prediction accuracy. Methods like FRASE-bot [9] apply deep learning to identify and score small molecular fragments that bind to different regions of the protein surface, which is often beneficial for the proteins without well-defined pockets [10].

Finally, the hybrid methods such as DELIA [11], PGpocket [12] and P2Rank [13] combine multiple predictive approaches to overcome limitations of individual techniques. However, this has a disadvantage of higher computational demands and large number of parameters.

Despite the large diversity of available techniques, most of them tend to identify more pocket-like regions than actually justified by experimental data. Nowadays it is generally straightforward for a researcher to find a bunch of cavities, pits and grooves on the surface of any protein that are potentially capable of binding to different kinds of ligands, while only some of them are functionally significant in the biological context. This redundancy of pocket detection is desirable in certain scenarios, such as identification of the previously unknown allosteric, hidden or “cryptic” binding pockets, but becomes a significant obstacle otherwise.

One of the most common tasks in computational drug discovery is docking the ligands into the known binding site of the target protein. In order to set up the docking procedure the researcher has to know the exact geometric extents of the binding pocket - something that is almost never explicitly annotated neither in the public databases nor in the research papers. The most obvious solution is to run one of the pocket prediction techniques, which provides such information. However, the result often includes redundant predicted pockets that partially overlap with the desired spatial or sequence region, so it is unclear which of them should be used as is, merged together or discarded. Correct post-processing of such predictions usually requires extensive analysis of the research papers, which is both time-consuming and error-prone. This issue becomes particularly pressing in the high-throughput or time-constrained scenarios in commercial drug discovery when human assessment of the binding pockets becomes the major bottleneck and a source of errors.

In this paper, we propose a hybrid approach that combines the geometry-based detection of the pockets with the Large Language Models, which assess them against the research papers and retain biologically relevant ones. Our intention is to test whether modern LLMs are powerful and accurate enough to substitute human researchers in binding pocket validation and prioritization tasks.

Specifically, we use Fpocket to generate multiple pocket candidates and LLMs to identify those that are supported by literature evidence. The LLMs are used to extract information from the literature about the residues associated with experimentally confirmed binding sites, which is then used to select the relevant pockets (Fig. 1).

**Figure 1.**
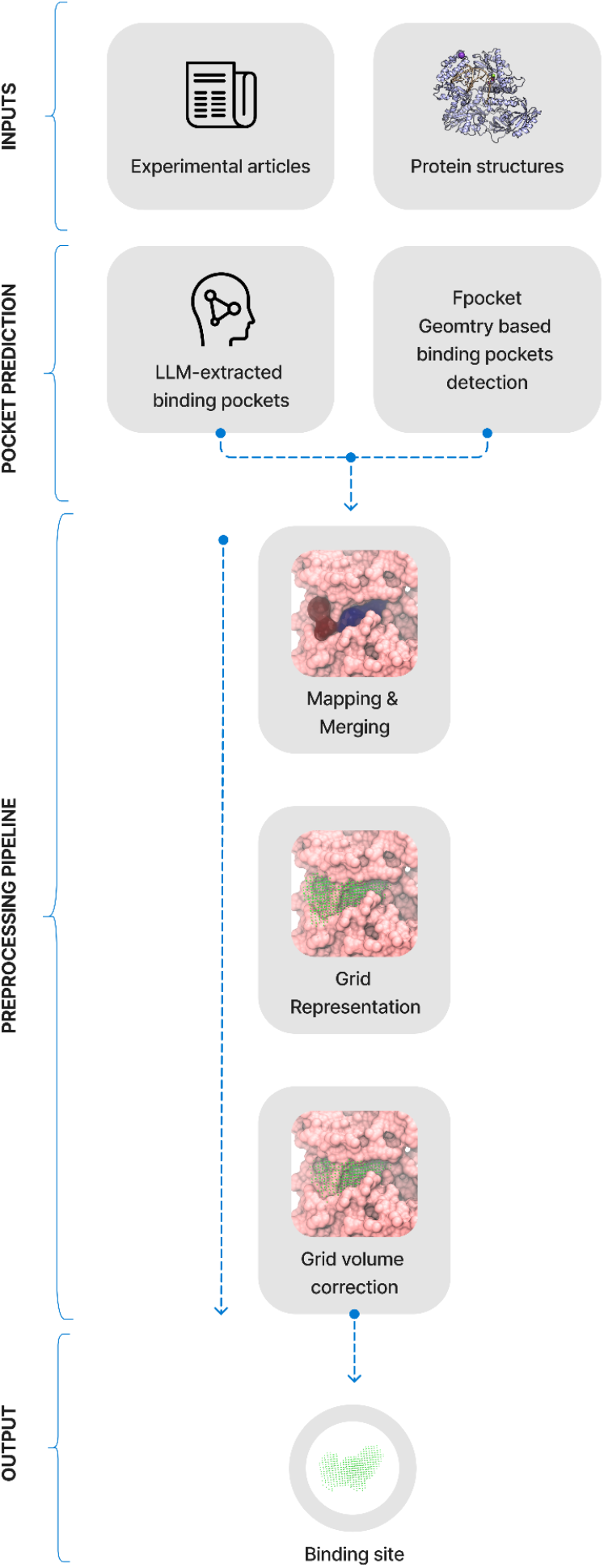
The overview of the proposed pipeline for the hybrid binding pocket detection. Fpocket – the geometry-based method – predicts numerous pocket candidates. The LLM analyses scientific articles and selects biologically relevant pockets. The results are then mapped and merged, so that LLM-extracted pockets are primarily used as filters for Fpocket predictions. Finally, the Fpocket pocket representation is converted into the spatial grid that undergoes postprocessing and refinement.

We have created an open-source benchmark dataset to evaluate LLM performance specifically for protein binding site detection tasks.

## 2. Methods

### 2.1. Dataset

At the time of writing there are no datasets for evaluating the performance of the protein binding pocket triaging and post-processing. To address this problem, we created a curated and manually annotated dataset that can be used as a public benchmark for LLM-based extraction of pocket-related information from the literature.

We selected ten diverse proteins: DNA polymerase alpha catalytic subunit; Tyrosine-protein kinase ABL1; 5-hydroxytryptamine receptor 2A; Muscarinic acetylcholine receptor M2; Voltage-gated sodium channel type 7; Voltage-gated sodium channel from from American cockroach; Gamma-aminobutyric acid receptor; GTPase KRas; Dihydroorotate dehydrogenase; Mixed lineage kinase domain-like protein.

For each of these targets, we found 2-6 peer-reviewed research articles (35 in total) and 3D structures from the Protein Data Bank (PDB). Selected articles represent different complexity tiers for extracting pocket-related information:

- Articles that do not describe binding sites are used as a negative control and allow false positives to be caught (11 articles).
- Articles that describe only one binding site for a given protein target. This is the easiest tier, which reveals LLM’s general understanding of structural biology concepts (17 articles).
- Articles that describe multiple binding pockets within a given target protein to evaluate the ability of LLM to precisely attribute specific amino acids to distinct pockets (3 articles).

The resulting benchmark dataset comprises JSON files, one for each selected paper. These files detail the small molecule binding sites explicitly described in the text, including the site’s name, description, and constituent amino acids.

### 2.2. LLM-based binding pocket extraction

In order to keep our technique reasonably fast and cost-effective we omitted advanced reasoning architectures like Chain of Thought (CoT) [14] or Tree of Thought (ToT) [15] — methods designed to guide complex, multi-step problem-solving. Instead, we relied on direct prompting, using examples and detailed instructions to steer the model. The final workflow incorporates three sequential steps:

1) paper relevance assessment, 2) pocket extraction and 3) pocket refinement, conducted by the same LLM model (Fig. 2). The full text of each article was processed in full since all papers fit into the context window of modern LLMs.

**Figure 2.**
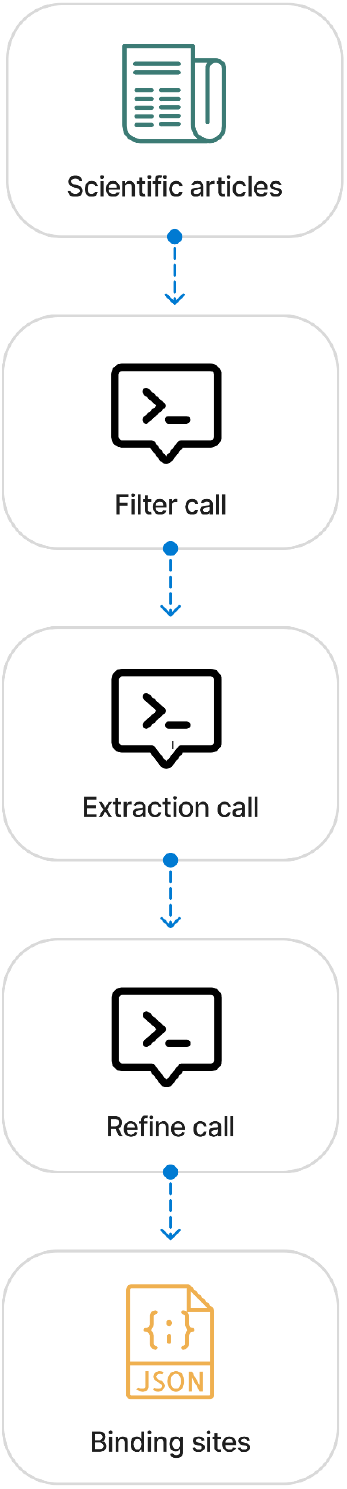
Overview of the LLM pocket extraction pipeline utilizing direct prompting. The pipeline processes complete articles through three LLM stages: (1) Filtering removes irrelevant papers lacking target relevance or residue-level descriptions. (2) Extraction, guided by extensive prompt optimization for format and accuracy, identifies small molecule binding site details (name, description, residues). (3) Refinement further improves data quality by correcting errors and omissions.

**Figure 3.**
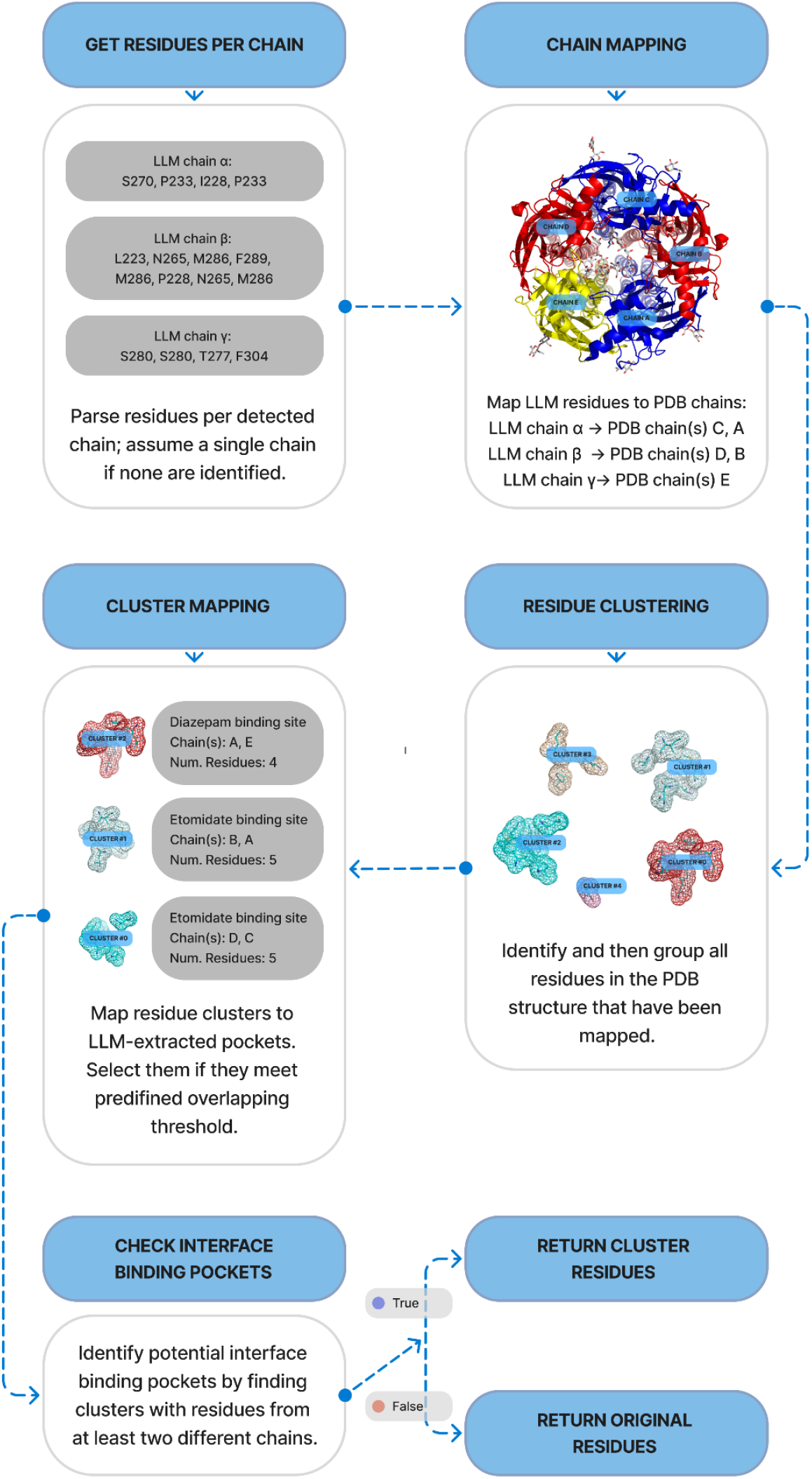
Workflow of the residue mapping algorithm. (1) LLM-extracted residues are assigned to potential PDB chains. (2) All mapped residues in the PDB structure undergo spatial clustering. (3) Resulting residue clusters are matched against LLM-defined pockets using an overlap threshold. (4) Matched clusters containing residues from multiple chains are identified as interface pockets and their residues are output. (5) Otherwise, original mapped residues are used.

In the first step we assess whether the provided paper is relevant for searching for the pockets for a given target protein i.e. whether it describes the correct protein and whether it contains description of the binding pockets at the residue level.

The pocket extraction step focuses on parsing the text and identifying amino acid residues comprising distinct binding pockets. Each pocket is assigned a name, description, and a list of residues in a predefined format. Significant prompt optimization efforts were focused on this step to ensure both strict output formatting and the correct identification of binding pockets.

Subsequent pocket refinement step evaluates the extraction results to correct any remaining inaccuracies such as identifying and including missing residues, rectifying improperly segmented binding sites, and filtering out binding sites irrelevant for small molecule binding (such as involved in PPI or DNA/RNA binding). Validation checks were performed during extraction and refinement specifically to guarantee the correct and consistent formatting of residue details (chain ID, residue name, and residue ID).

### 2.3 Pocket extraction accuracy metrics

To assess the performance of our pipeline, we calculated the following pocket- and paper-based metrics:

#### Pocket Number Accuracy

for each paper, if the number of annotated and extracted pockets match, then prediction is correct (1), otherwise it is incorrect (0). Based on these values for each pocket the final accuracy per parer is calculated.

#### Pocket Recall

represents the percentage of annotated pockets that have been correctly extracted by the LLM:

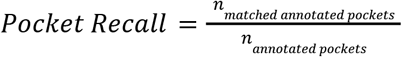

#### Pocket Specificity (True Negative Rate, TNR)

Reflects how often the pipeline correctly avoids predicting a pocket when none is annotated in the ground truth. This is calculated as 1 - False Positive Rate (FPR). The FPR indicates the frequency at which the pipeline generates a fake pocket that does not correspond to any actual annotated pocket.

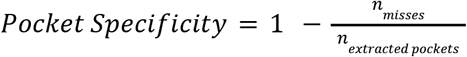

where *n*_*misses*_ − the number of parsed pockets with no correspondence to any annotated pocket.

To evaluate performance at the amino acid level within the pockets, we track the F1 score, recall, and precision metrics for the lists of amino acids in annotated and extracted pockets. The primary optimization objectives for our pipeline are maximizing the F1 score at the amino acid level, minimizing the Pocket False Positive Rate (FPR), and maximizing Pocket Accuracy and Pocket Recall. While other metrics are monitored to evaluate overall pocket extraction quality, they do not guide the optimization process.

### 2.4 Pocket clustering and post-processing

The list of amino acids extracted by the LLM from the papers may not map directly into a specific PDB structure due to the sequence differences (e.g., insertions/deletions and engineered mutations) or errors in residue numbers, names or chain IDs of the LLM extracted residues. Therefore, an additional algorithmic mapping step is required to mitigate these issues. It allows handling such cases as assigning residues consistently across identical chains in homomers and correctly identifying binding sites situated at the interfaces between different chains, which are otherwise very hard to address on the level of LLM extraction.

First, chain correspondence between the paper’s residue descriptions and the specific PDB structure is established using a sequence similarity threshold. Following this, spatial clustering is performed on all identified pocket residues within the PDB to detect potential inter-chain interfaces. A key step involves determining if these spatially clustered inter-chain residue groups match any specific pockets extracted by the LLM using a different sequence similarity threshold. If a match is found, the amino acids from that corresponding cluster are used, defining a unique inter-chain pocket. If no such match is identified, the LLM-extracted residues for the pocket are instead mapped onto all potentially relevant PDB chain identifiers, indicating the possibility of multiple, non-unique pockets located on different protein subunits.

Although clustering can be applied to identify all binding sites, we found this approach yields suboptimal results when distinct binding sites are located in close spatial proximity. However, clustering is particularly effective at capturing identical binding sites formed at the interfaces between subunits in homo- or hetero-multimers.

Manipulations with protein structures were performed with the custom scripts based on MolAR molecular modeling library [16].

### 2.5 Construction of pocket volumetric representation

Since residue information alone lacks the spatial detail to define a volumetric binding pocket, a geometry-based method provides this necessary geometric representation, while the LLM output subsequently serves as a filter to select the most plausible candidates among these geometric predictions. For geometry-based prediction, we employ Fpocket, a method that identifies potential binding sites using Voronoi tessellation and alpha spheres. These predictions are subsequently filtered based on the LLM output, retaining only those pockets whose alpha spheres are located near the LLM extracted residues.

If multiple LLM-extracted pockets overlap with the same geometric pocket, they are merged into a single pocked if the residue overlap between them exceeds a predefined threshold. This also allows blending the data from different publications describing the same binding pocket on the target protein, but report different or incomplete sets of associated residues.

The space bounded by the binding pocket surface can be filled with either alpha-spheres or grid points depending on the used technique. Multiple geometry-based approaches such as CB-Dock2 [17], CASTp 3.0 [18], PyVOL [19] and Fpocket itself [1] use spheres to represent protein cavities. Grid representation is used by DoGSite3 [20] and POVME 2.0 [21].

The representation of alpha spheres used by Fpocket is not very practical for our purposes because merging or splitting the pockets represented by spheres is non-trivial and involves unnecessarily complex algorithms. Thus, we convert the pockets to a grid representation, as shown in Figure 4.

**Figure 4.**
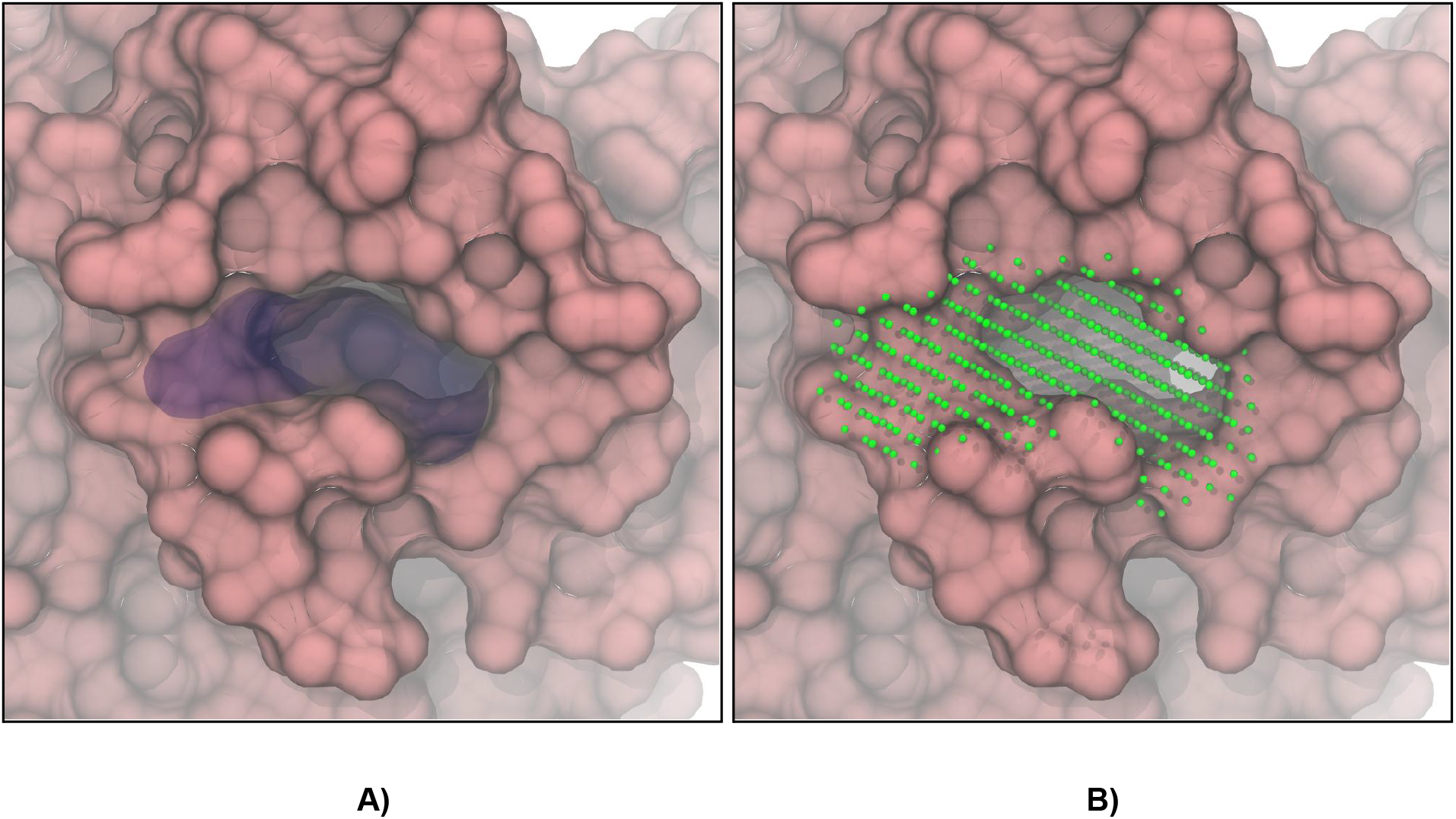
Mapped and filtered pockets for the inhibitor site of Dihydroorotate dehydrogenase (PDB: 6OC0) visualized using A) Fpocket alpha spheres representation; B) grid representation.

We define a rectangular grid with 1.5Å steps whose boundaries correspond to the minimum and maximum coordinates of the binding site heavy atoms along each axis. Only grid points located inside the alpha spheres are retained.

The alpha spheres may protrude significantly from the protein cavities into the bulk solvent. The grid representations of such spheres overestimate the pocket volume and may lead to artifacts if this volume is used for molecular docking by allowing much bulkier compounds than can actually fit into the pocket. To address this, we construct a convex hull on the pocket heavy atoms and remove all grid points that fall outside it (Fig. 5).

**Figure 5.**
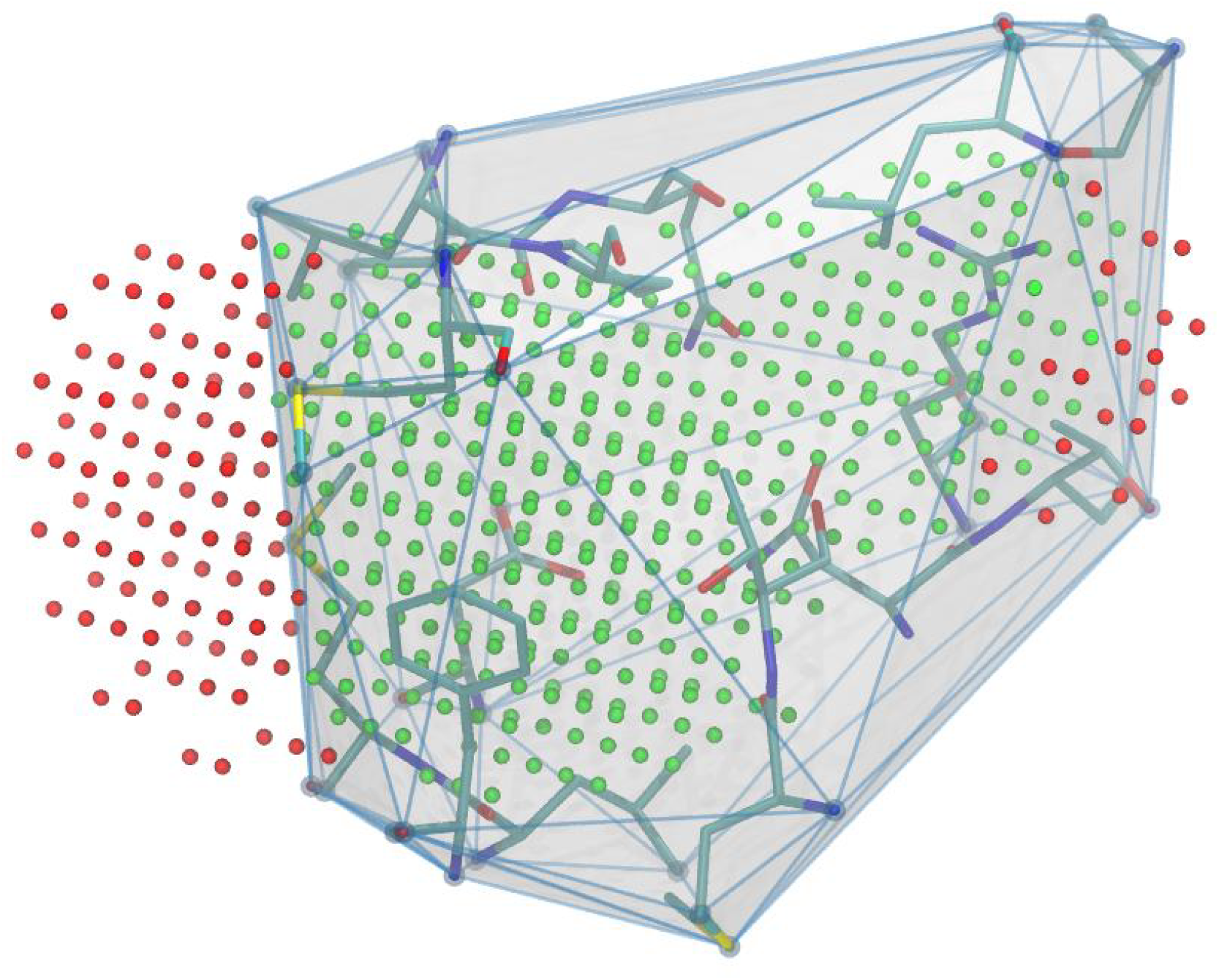
Filtering pocket grid points using a convex hull defined by pocket atoms. Grid points falling outside the hull boundary (red) are discarded, while those inside (green) are retained, resulting in a corrected pocket representation without protruding parts.

**Figure 6.**
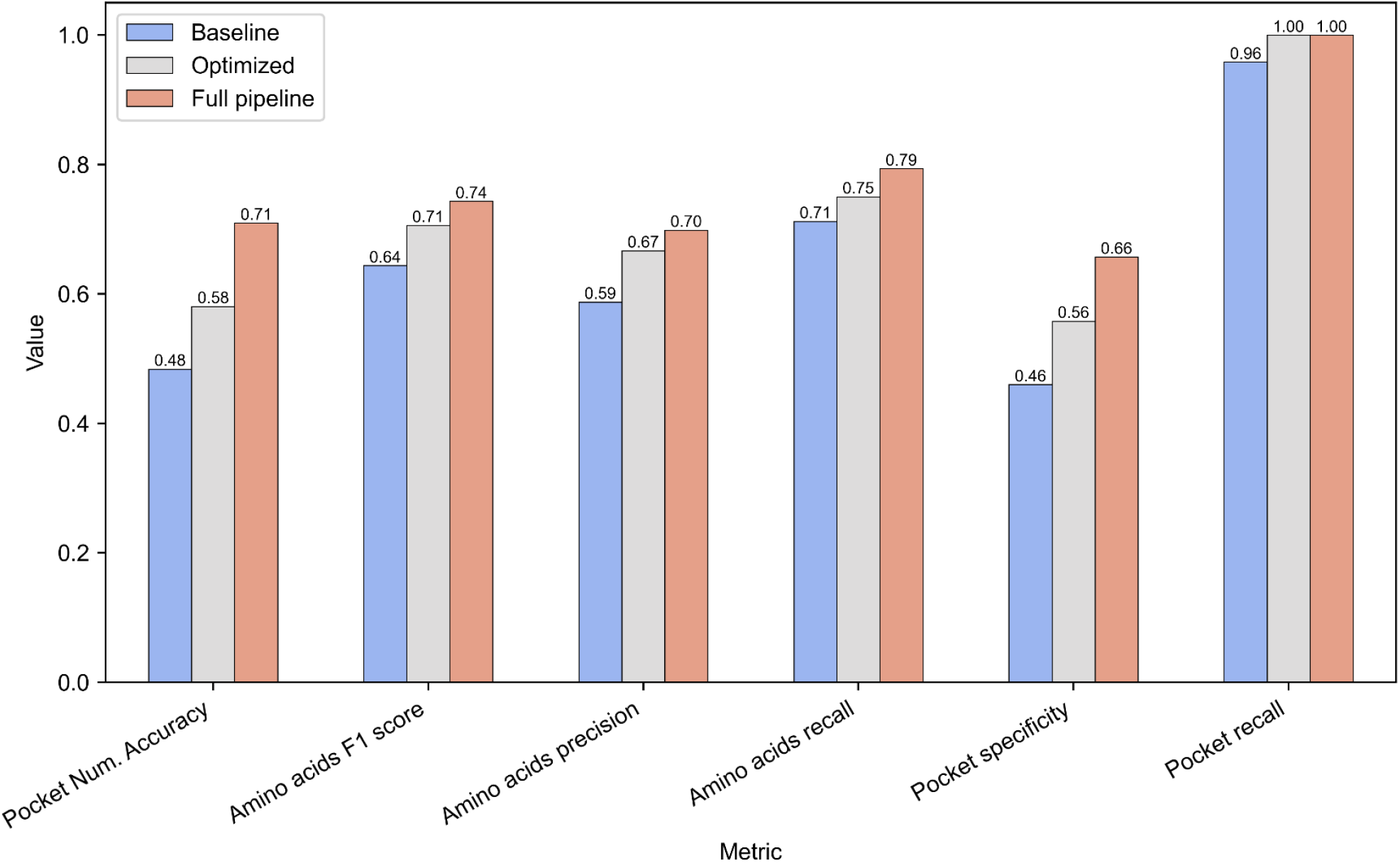
GPT-4o-mini performance metrics for three pipeline configurations: Baseline (filter and extraction steps with initial prompts), Optimized (filter and extraction steps with optimized prompts), and Full pipeline (adding a refinement step to the Optimized configuration).

The resulting grid is further refined by deleting the points located closer than the Van der Waals radius to any protein atom. This step is not strictly necessary since the docking algorithms prevent the sterical slashes anyway, but it improves the visual quality of the pocket representation.

## 3. Results

### 3.1 LLM pipeline performance

Due to the rapid advancement of LLMs, our development and testing focused exclusively on the model versions actual at the time of writing, and results are reported accordingly. Our goal is to show that a reliable pocket prioritization pipeline can be built with the LLM technologies at the time of writing. All steps of the LLM pipeline are performed by the GPT-4o-mini model.

The dataset was initially evaluated by the baseline set of prompts (see Supplementary Information). Then the prompts for filtering and extractions stages were optimized using the following strategies:

- Adding explicit instructions to ensure that LLM identifies only binding sites relevant to small molecules, while actively excluding other functional regions such as protein-protein interaction (PPI) sites and areas involved in DNA/RNA binding.
- Adding directives to only include amino acids explicitly involved in pocket formation or ligand interaction.
- Detailed specifications of the response format and quality standards, including guidelines that strongly rewards correct residue reporting while penalizing hallucinated or inaccurate entries.
- Enforcement of strict formatting rules during residue extraction, ensuring the correct parsing and consistent representation of chain identifiers, residue names, and residue IDs.

These changes were implemented sequentially with monitoring performance following each prompt adjustment. Adjustments were only retained if they improved the metrics and/or correctness of output formatting. The final prompts are available in Supplementary Information.

The initial Baseline configuration utilized non-optimized prompts for extraction and filtering. This yielded a Pocket Number Accuracy of 0.484, Pocket Recall of 0.958, and Pocket Specificity of 0.46.

Optimization of the extraction step (while keeping the filtering unchanged) boosted performance significantly. It resulted in a Pocket Number Accuracy of 0.581, perfect Pocket Recall (1.0), and Pocket Specificity of 0.558. However, manual inspection revealed that this configuration sometimes incorrectly split single pockets into multiple entries.

Addition of the Refinement step addressed this issue, while further improving Pocket Number Accuracy to 0.71 and Pocket Specificity to 0.657 and maintaining perfect Pocket Recall. This final step also enhanced residue-level metrics, as the refinement process often incorporates missing amino acids.

The filtering step was also evaluated separately as a binary classification task to determine paper relevance based on the presence of binding site descriptions. The model achieved an accuracy of 0.87, recall of 1.0, precision of 0.833, and a False Positive Rate (FPR) of 0.363, while retaining FNR at 0. The perfect recall indicates the model effectively identifies all of the relevant papers. However, the precision and FPR suggest that approximately 36% of papers flagged as relevant are actually false positives that do not get filtered out at this stage. We considered this acceptable because inclusion of potentially irrelevant papers is unlikely to predict fake pockets that are hard to spot on the post-processing.

### 3.2 Volumetric pocket representations

We visually assessed the residue mapping algorithm’s performance for each target structure, with the quality of the final mapping assessed manually by a human expert. This assessment focused primarily on ensuring correct residue assignments across all relevant PDB chains, as these would influence the pocket’s volumetric representation. To achieve optimal performance across the whole dataset, we empirically set the threshold 0.6 for chain matching and 0.7 for cluster matching. Figure 7 shows mapped residues for the GABAA receptor PDB structure, where the residue mapping algorithm identified two equivalent binding pockets at distinct subunit interfaces. The output includes atom coordinates for the pocket’s constituent residues along with the corresponding pocket name and description extracted by the LLM.

**Figure 7.**
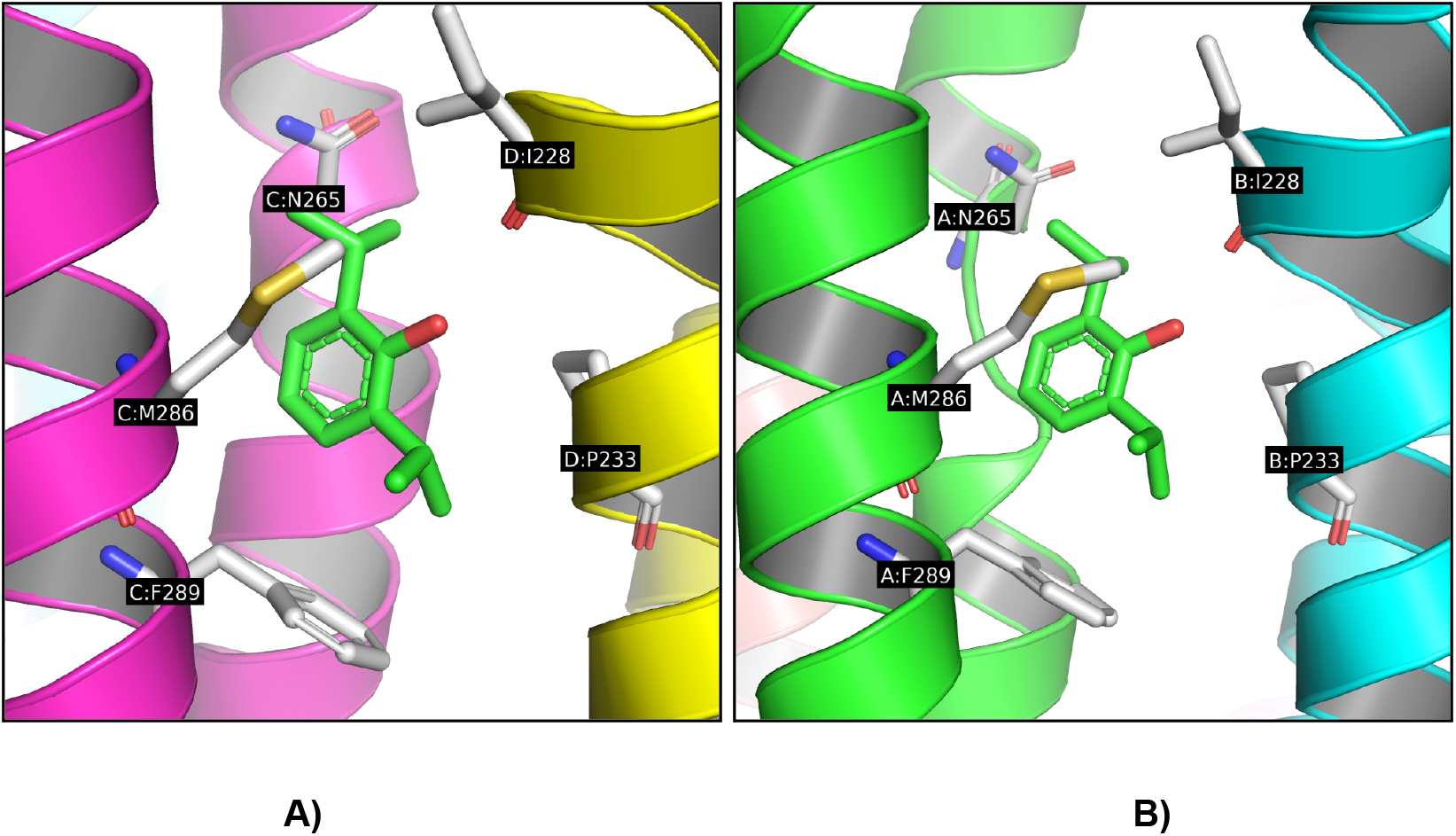
Proposed algorithm’s residue mapping showing equivalent etomidate binding pockets in the GABAA receptor (PDB: 6X3T) at (A) the A:B interface and (B) the C:D interface [22].

We generated initial geometric pocket representations using Fpocket with default parameters. However, because Fpocket sometimes fragments larger sites, a post-processing step is implemented to merge spatially adjacent pockets meeting a predefined empirical residue overlap threshold. Kinase structures frequently necessitate such merging, as Fpocket often splits their ATP-binding sites into two or more smaller subpockets that must be combined to accurately represent the complete catalytic site (Fig. 8).

**Figure 8.**
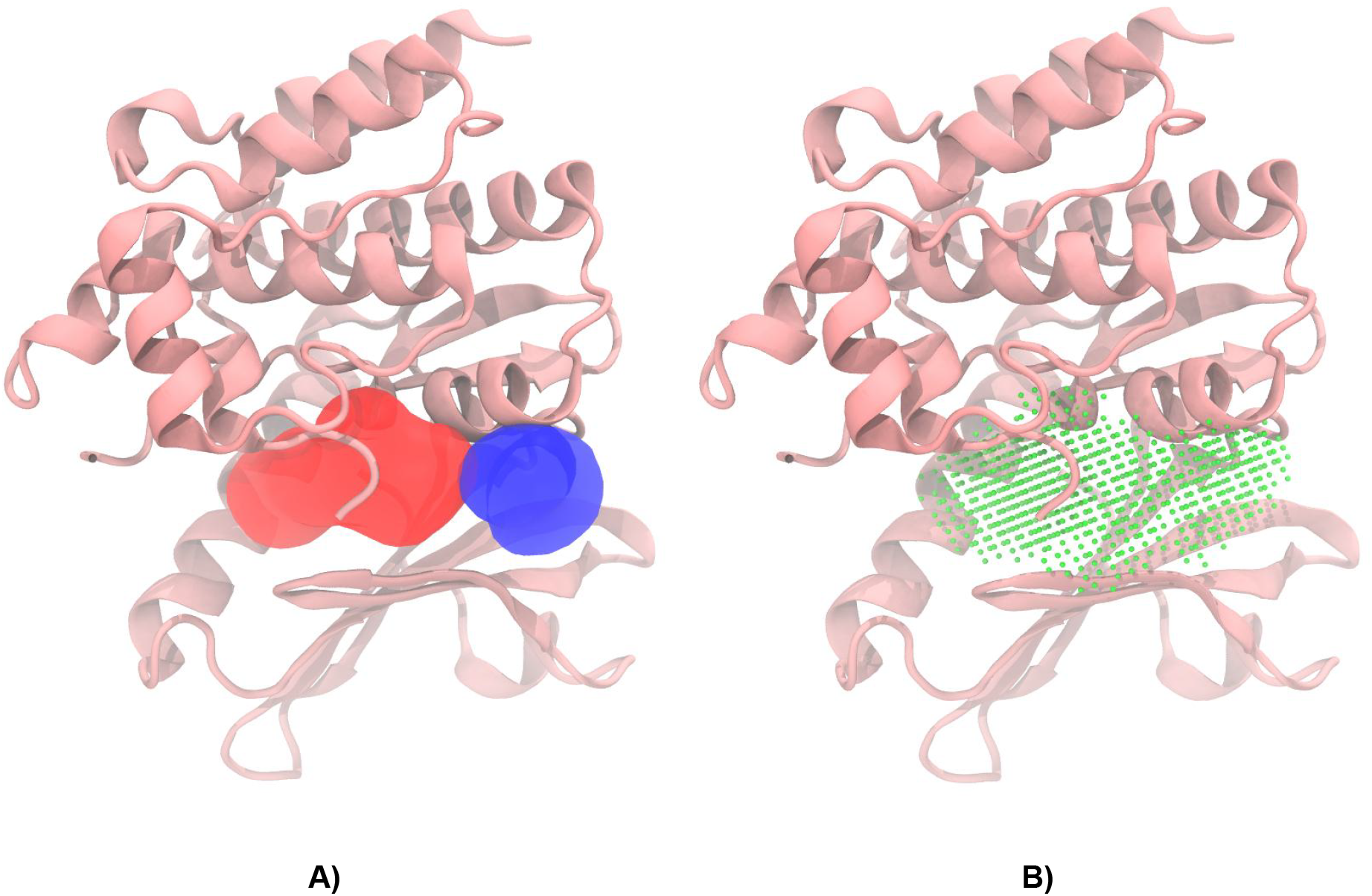
Mixed lineage kinase domain like pseudokinase (PDB: 4MWI) with (A) selected pockets as alpha spheres and (B) the final pocket after merging and post-processing [23].

The same merging strategy addresses another challenge: consolidating information from multiple papers. Pockets identified from different sources might describe the same physical site but often list different constituent residues. The sequence-level comparisons could be attempted, however merging based on the spatial proximity of the pockets is a more robust approach for identifying and unifying equivalent sites across various publications.

This merging process, however, cannot address the cases where Fpocket initially fails to separate distinct binding sites and outputs them incorrectly as a large single pocket. This limitation was observed with the M2 receptor, where Fpocket did not distinguish between known separate binding sites (Fig. 9). Despite these challenges, the residue mapping algorithm performed reliably for binding sites contained within a single chain or distinct protein domain.

**Figure 9.**
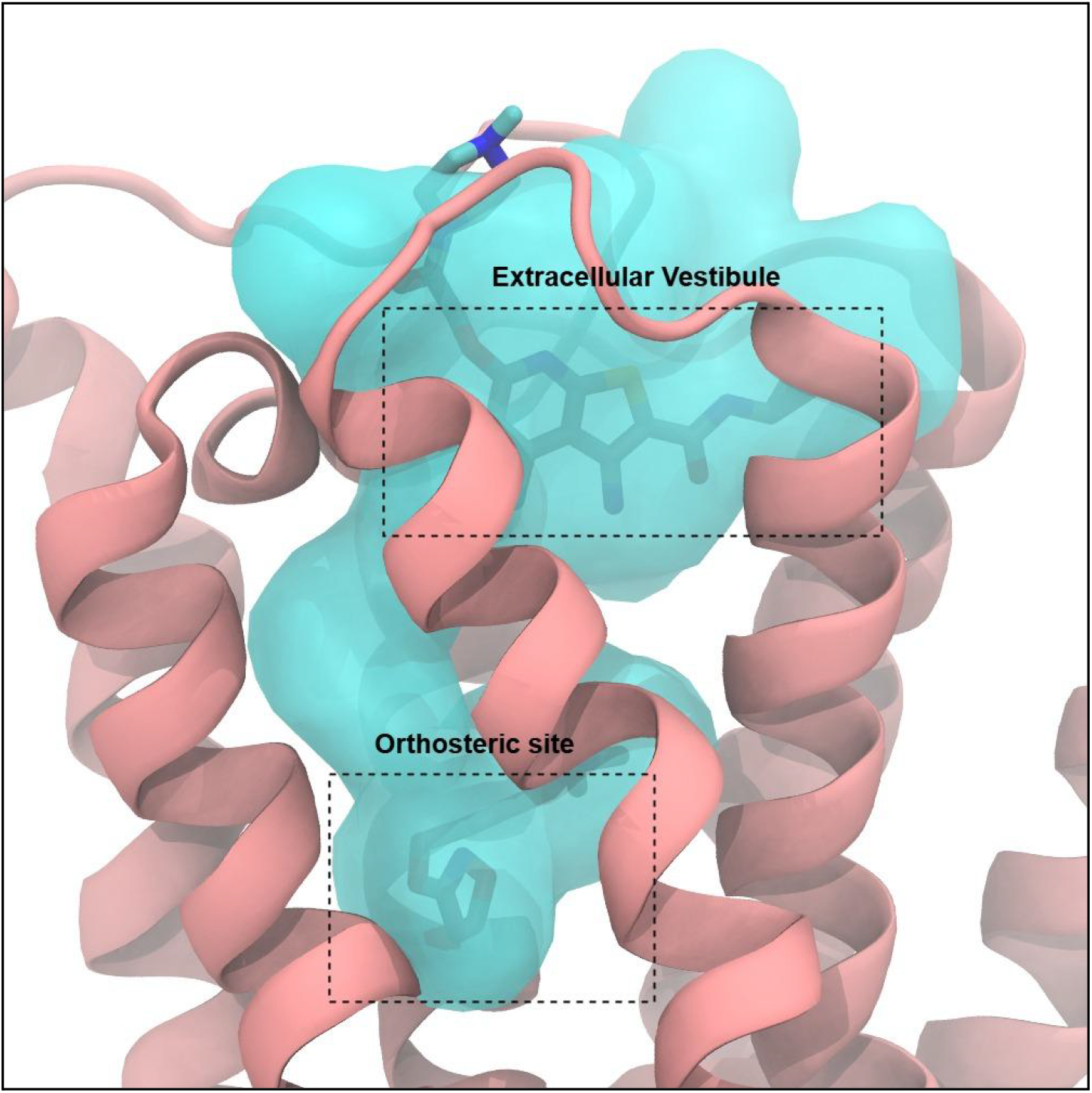
Muscarinic acetylcholine receptor M2 (PDB: 4MQT) with detected and mapped pocket that combines the orthosteric and extracellular vestibule binding sites [24].

For multimeric assemblies the merging algorithm only performs reliably assuming correct initial residue identification by the LLM. For instance, the human Nav1.7-VSD4-NavAb chimera (PDB: 5EK0) is a tetramer complex where each of four chains incorporates a human Na_V_1.7 VSD4 domain introduced via protein engineering. In this case, our algorithm correctly located four identical binding pockets — one within each of the VSD4 domains (Figure 10) that correspond to the known deep binding site of the GX-936 warhead in the VSD4 extracellular cleft [25].

**Figure 10.**
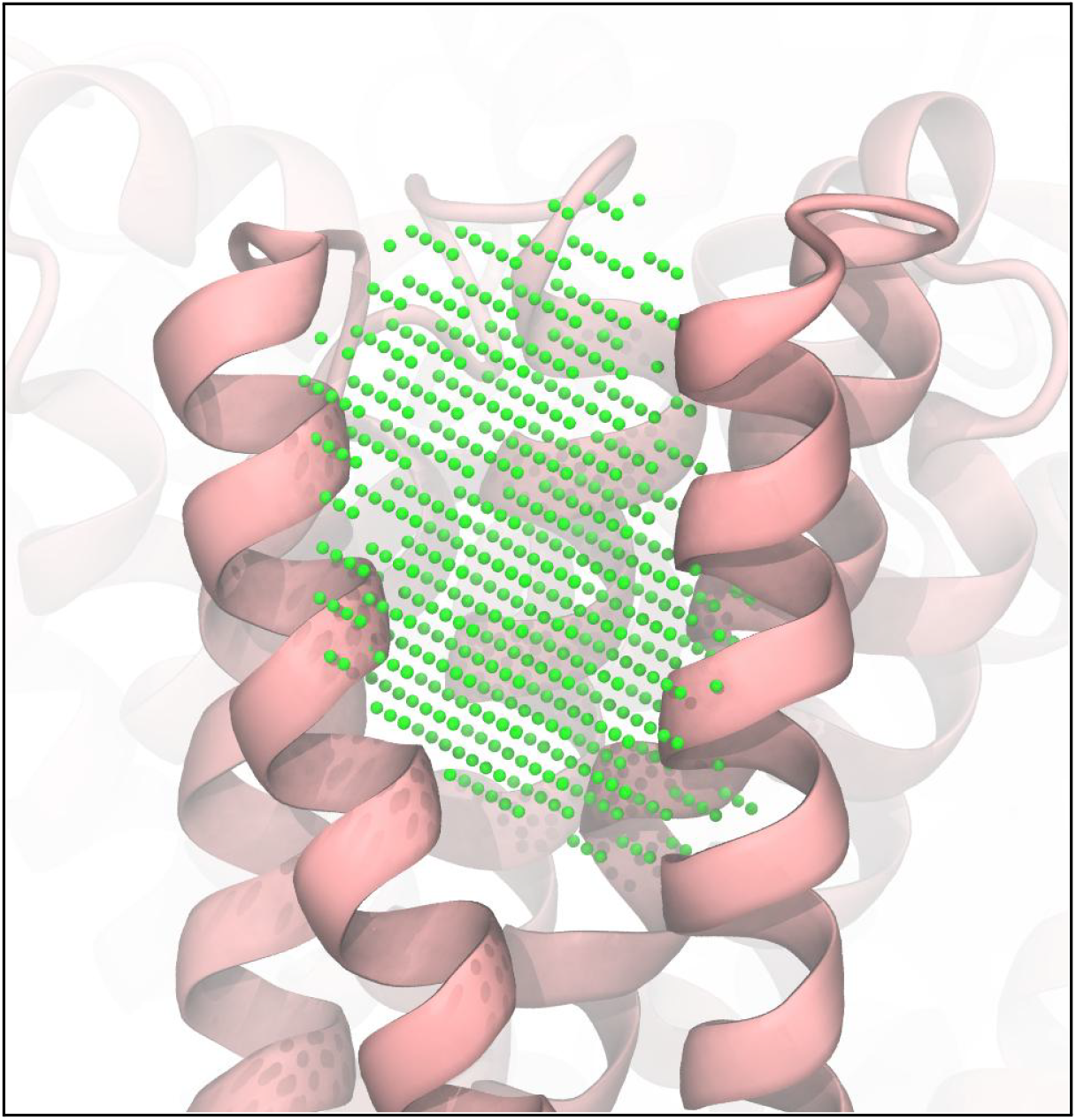
GX-936 binding pocket located at the VSD4 domain of the human Na_V_1.7 (PDB: 5EK0).

## 4. Discussion

The identification and prioritization of protein binding pockets is a critical step in small molecule drug discovery. While numerous computational methods exist, practical workflows often face challenges, particularly when focusing on biologically validated binding sites relevant for virtual screening or druggability assessment, rather than discovering entirely novel pockets. Selecting and defining these known sites typically involves a laborious process: manually analyzing multiple research papers to identify key residues, dealing with inconsistent terminologies and data presentation across publications, computing volumetric representations (which are rarely provided directly), and mapping the literature-derived residue information onto these geometric pockets to filter for biological relevance. This manual workflow is time-consuming and prone to errors, especially in high-throughput industrial settings.

This paper investigates the potential of automating this workflow using a hybrid approach combining geometry-based pocket detection with Large Language Models (LLMs) for literature analysis and pocket validation. We aimed to determine if current LLM technology can effectively substitute human researchers in the binding pocket validation and prioritization tasks outlined above. Our proposed pipeline integrates Fpocket for initial geometric pocket prediction with an LLM (specifically, GPT-4o-mini) for extracting and validating binding site information from scientific literature, followed by algorithmic mapping, merging, and refinement steps to produce accurate volumetric pocket representations.

Since parsing complex PDF files of scientific articles remains challenging at the time of writing, we prepared manually curated markdown representations of the articles for this study. In practice the quality of the parsed textual representation of the article may significantly restrict the model performance. This is especially important for information contained in tables and figures, which is notoriously difficult to parse correctly. Solving these issues is beyond the scope of the current work.

Our results indicate that the LLM pipeline, particularly after prompt optimization and the inclusion of a refinement step, can effectively identify relevant papers and extract pocket information with high accuracy. The refinement stage proved valuable in correcting initial extraction errors, such as improper pocket splitting or missing residues. However, some challenges remain: the LLM filtering step still passes some irrelevant papers through, and accurately interpreting the nuances of binding site descriptions spread across varied writing styles and sections within papers requires careful handling. Explicit instructions were needed to ensure that LLM focused only on small-molecule binding sites.

The subsequent algorithmic mapping and clustering successfully bridged the gap between literature descriptions and specific PDB structures, handling inconsistencies like sequence differences, numbering errors, and multimeric interfaces. This step demonstrated its utility in identifying equivalent pockets across different subunits, as seen in the GABAA receptor and Nav1.7 chimera examples. Nevertheless, its accuracy depends on the initial LLM extraction quality and the structural similarity between the PDB file and the experimental construct described. Multimeric proteins remain challenging in this regard due to possible confusion of residue attribution between the chains and positioning of the pockets between multiple subunits.

Generating the final volumetric representation involves filtering Fpocket’s geometric predictions using LLM-validated residues. Converting to a grid representation and subsequent post-processing, including merging fragmented pockets and applying convex hull truncation, significantly improved the final pocket definition. This merging also effectively consolidated information from multiple publications. Still, the pipeline inherits Fpocket’s limitations, such as its occasional inability to separate closely adjacent sites, as observed with the M2 receptor.

Despite existing limitations, including the reliance on the specific LLM used (GPT-4o-mini) and the limited benchmark dataset’s scope, our hybrid approach shows considerable promise. Future work could involve exploring more advanced LLM techniques, integrating alternative geometric predictors, expanding the benchmark dataset, and developing methods to automatically resolve structural discrepancies. Overall, leveraging LLMs to automate the analysis of research literature for binding pocket identification appears to be a valuable direction, potentially streamlining the drug discovery workflows.

## 5. Conclusion

This study demonstrates a promising new hybrid approach combining geometric pocket detection with LLM-based literature analysis to improve the identification and prioritization of biologically relevant protein binding pockets in the context of small molecule drug discovery. By leveraging the power of LLMs to extract experimental evidence from research papers, the method effectively filters out irrelevant geometric pocket predictions, offering a more accurate and efficient workflow for applications in the high-throughput automated drug discovery workflows. The benchmark dataset is made publicly available to further support future advancements in this field.

## 6. Author contributions

AN served as the conceptual lead and originator of the core idea. MM carried out the computational experiments, prepared datasets, wrote the analysis scripts and prompts, ran the tests and analysed results. ZO performed initial dataset preparation, wrote the code for LLM invocation and assembled the initial pipeline. IK, RS, VH, VB and NS participated in benchmark dataset preparation and validation. SY coordinated the work, critically assessed the results, guided the analysis and participated in prompt optimization. SS and AN designed the study and controlled its progress. The manuscript was written by MM and SY.

## 8. Acknowledgements

S.Y. has received funding through the grant MSMT-355/2025-16 from the Ministry of Education, Youth and Sport of the Czech Republic and through the MSCA4Ukraine project 101101923, which is funded by the European Union. Views and opinions expressed are however those of the author(s) only and do not necessarily reflect those of the European Union. Neither the European Union nor the MSCA4Ukraine Consortium as a whole nor any individual member institutions of the MSCA4Ukraine Consortium can be held responsible for them.

The authors acknowledge the use of the large language models GPT4o and Gemini 2.5 Pro for proofreading and stylistic editing of the manuscript.

## References

[1] V. Le Guilloux, P. Schmidtke, and P. Tuffery, “Fpocket: An open source platform for ligand pocket detection,” BMC Bioinformatics, vol. 10, no. 1, p. 168, Jun. 2009, doi: 10.1186/1471-2105-10-168.

[2] J. Liang, H. Edelsbrunner, and C. Woodward, “Anatomy of protein pockets and cavities: measurement of binding site geometry and implications for ligand design,” Protein Sci. Publ. Protein Soc., vol. 7, no. 9, pp. 1884–1897, Sep. 1998, doi: 10.1002/pro.5560070905.

[3] D. Ghersi and R. Sanchez, “Improving accuracy and efficiency of blind protein-ligand docking by focusing on predicted binding sites,” Proteins Struct. Funct. Bioinforma., vol. 74, no. 2, pp. 417–424, Feb. 2009, doi: 10.1002/prot.22154.

[4] J. A. Capra, R. A. Laskowski, J. M. Thornton, M. Singh, and T. A. Funkhouser, “Predicting Protein Ligand Binding Sites by Combining Evolutionary Sequence Conservation and 3D Structure,” PLOS Comput. Biol., vol. 5, no. 12, p. e1000585, Dec. 2009, doi: 10.1371/journal.pcbi.1000585.

[5] J. Jiménez, S. Doerr, G. Martínez-Rosell, A. S. Rose, and G. De Fabritiis, “DeepSite: protein-binding site predictor using 3D-convolutional neural networks,” Bioinformatics, vol. 33, no. 19, pp. 3036–3042, Oct. 2017, doi: 10.1093/bioinformatics/btx350.

[6] R. Aggarwal, A. Gupta, V. Chelur, C. V. Jawahar, and U. D. Priyakumar, “DeepPocket: Ligand Binding Site Detection and Segmentation using 3D Convolutional Neural Networks,” J. Chem. Inf. Model., vol. 62, no. 21, pp. 5069–5079, Nov. 2022, doi: 10.1021/acs.jcim.1c00799.

[7] J. Kandel, H. Tayara, and K. T. Chong, “PUResNet: prediction of protein-ligand binding sites using deep residual neural network,” J. Cheminformatics, vol. 13, no. 1, p. 65, Sep. 2021, doi: 10.1186/s13321-021-00547-7.

[8] K. Jeevan, S. Palistha, H. Tayara, and K. T. Chong, “PUResNetV2.0: a deep learning model leveraging sparse representation for improved ligand binding site prediction,” J. Cheminformatics, vol. 16, no. 1, p. 66, Jun. 2024, doi: 10.1186/s13321-024-00865-6.

[9] Y. An et al., “In silico fragment-based discovery of CIB1-directed anti-tumor agents by FRASE-bot,” Nat. Commun., vol. 15, no. 1, p. 5564, Jul. 2024, doi: 10.1038/s41467-024-49892-9.

[10] Y. An et al., “Machine Learning-driven Fragment-based Discovery of CIB1-directed Anti-Tumor Agents by FRASE-bot,” Res. Sq., p. rs.3.rs-3197490, Aug. 2023, doi: 10.21203/rs.3.rs-3197490/v1.

[11] C.-Q. Xia, X. Pan, and H.-B. Shen, “Protein-ligand binding residue prediction enhancement through hybrid deep heterogeneous learning of sequence and structure data,” Bioinforma. Oxf. Engl., vol. 36, no. 10, pp. 3018–3027, May 2020, doi: 10.1093/bioinformatics/btaa110.

[12] Y. Zhao et al., “A Point Cloud Graph Neural Network for Protein-Ligand Binding Site Prediction,” Int. J. Mol. Sci., vol. 25, no. 17, p. 9280, Aug. 2024, doi: 10.3390/ijms25179280.

[13] R. Krivák and D. Hoksza, “P2Rank: machine learning based tool for rapid and accurate prediction of ligand binding sites from protein structure,” J. Cheminformatics, vol. 10, no. 1, p. 39, Aug. 2018, doi: 10.1186/s13321-018-0285-8.

[14] J. Wei et al., “Chain-of-Thought Prompting Elicits Reasoning in Large Language Models,” Jan. 10, 2023, arXiv: arXiv:2201.11903. doi: 10.48550/arXiv.2201.11903.

[15] S. Yao et al., “Tree of Thoughts: Deliberate Problem Solving with Large Language Models,” Dec. 03, 2023, arXiv: arXiv:2305.10601. doi: 10.48550/arXiv.2305.10601.

[16] S. Yesylevskyy, “MolAR: Memory-Safe Library for Analysis of MD Simulations Written in Rust,” J. Comput. Chem., vol. 46, no. 1, p. e27536, 2025.

[17] Y. Liu, X. Yang, J. Gan, S. Chen, Z.-X. Xiao, and Y. Cao, “CB-Dock2: improved protein-ligand blind docking by integrating cavity detection, docking and homologous template fitting,” Nucleic Acids Res., vol. 50, no. W1, pp. W159–W164, Jul. 2022, doi: 10.1093/nar/gkac394.

[18] W. Tian, C. Chen, X. Lei, J. Zhao, and J. Liang, “CASTp 3.0: computed atlas of surface topography of proteins,” Nucleic Acids Res., vol. 46, no. W1, pp. W363–W367, Jul. 2018, doi: 10.1093/nar/gky473.

[19] “PyVOL Documentation — PyVOL 1.7.6 documentation.” Accessed: Apr. 24, 2025. [Online]. Available: https://schlessinger-lab.github.io/pyvol/

[20] J. Graef, C. Ehrt, and M. Rarey, “Binding Site Detection Remastered: Enabling Fast, Robust, and Reliable Binding Site Detection and Descriptor Calculation with DoGSite3,” J. Chem. Inf. Model., vol. 63, no. 10, pp. 3128–3137, May 2023, doi: 10.1021/acs.jcim.3c00336.

[21] J. D. Durrant, L. Votapka, J. Sørensen, and R. E. Amaro, “POVME 2.0: An Enhanced Tool for Determining Pocket Shape and Volume Characteristics,” J. Chem. Theory Comput., vol. 10, no. 11, pp. 5047–5056, Nov. 2014, doi: 10.1021/ct500381c.

[22] J. J. Kim et al., “Shared structural mechanisms of general anaesthetics and benzodiazepines,” Nature, vol. 585, no. 7824, pp. 303–308, Sep. 2020, doi: 10.1038/s41586-020-2654-5.

[23] J. M. Murphy et al., “Insights into the evolution of divergent nucleotide-binding mechanisms among pseudokinases revealed by crystal structures of human and mouse MLKL,” Biochem. J., vol. 457, no. 3, pp. 369–377, Feb. 2014, doi: 10.1042/BJ20131270.

[24] A. C. Kruse, B. K. Kobilka, D. Gautam, P. M. Sexton, A. Christopoulos, and J. Wess, “Muscarinic acetylcholine receptors: novel opportunities for drug development,” Nat. Rev. Drug Discov., vol. 13, no. 7, pp. 549–560, Jul. 2014, doi: 10.1038/nrd4295.

[25] S. Ahuja et al., “Structural basis of Nav1.7 inhibition by an isoform-selective small-molecule antagonist,” Science, vol. 350, no. 6267, p. aac5464, Dec. 2015, doi: 10.1126/science.aac5464.

