## Supplementary Information for "Leveraging Large Language Models for Literature-Driven Prioritization of Protein Binding Pockets"

*Mykola Melnychenko* [1], *Semen Yesylevskyy* [1,2,3,4], Zakhar Osrovsky [1], *Serhii Starosyla* [1], *Ihor Koleiev* [1,3], *Taras Voitsitskyi* [1,3], *Roman Stratiichuk* [1,5], *Vladyslav Husak* [1,7], *Volodymyr Bdzhola* [6], *Nazar Shevchuk* [1], *Alan Nafiiev* [1].

[1] - Receptor.AI Inc., 20-22 Wenlock Road, London N1 7GU, United Kingdom.
[2] - Institute of Organic Chemistry and Biochemistry, Czech Academy of Sciences, CZ-166 10 Prague 6, Czech Republic.
[3] - Department of Physics of Biological Systems, Institute of Physics of The National Academy of Sciences of Ukraine, 46 Nauky Ave., 03038, Kyiv, Ukraine.
[4] - Department of Physical Chemistry, Faculty of Science, Palacký University Olomouc, 17. listopadu 12, 771 46 Olomouc, Czech Republic.
[5] - Department of Biophysics and Medical Informatics, Educational and Scientific Centre "Institute of Biology and Medicine", Taras Shevchenko Kyiv National University, 64 Volodymyrska Str., 01601, Kyiv, Ukraine.
[6] - Institute of Molecular Biology and Genetics of The National Academy of Sciences of Ukraine, 150 Zabolotnogo Str., 03143, Kyiv, Ukraine
[7] -Department of Cellular, Computational and Integrative Biology, The University of Trento, Via Sommarive 9, 38123 Povo (Trento), Italy

| Model | Prompts |
|---|---|
| Baseline | There is a scientific paper text to analyze indicated by <article> html tags. Do the following:<br>1. Determine the number of unique binding sites for small molecules in target protein "{target_protein}" that are described in the text.<br>2. Provide a very laconic, very specific and discriminative characteristic for each binding site that you identified.<br>3. If the context permits, include any relevant small molecules in the binding site description.<br>4. Output the list of amino acid residues constituting each of identified binding sites. |

<table>
<tr>
<td></td>
<td>Use the following notation for defining amino acid residues:<br><chain_id><res_name><res_num>.<br>4.1 chain_id is an optional chain identifier for each amino acid.<br>4.2 res_name is the amino acid name in either single-letter or three letter notation. Use the same notation as in the paper.<br>4.3 res_num is the integer amino acid number.<br>4.4 Some residues may be followed by a punctuation mark or a space. Any characters that follow them are not part of the residue.</td>
</tr>
<tr>
<td>Optimized</td>
<td>There is a scientific paper text to analyze indicated by <article> html tags. Do the following:<br>1. Determine the number of unique binding sites for small molecules in target protein "{target_protein}" that are described in the text.<br>2. Provide a very laconic, very specific and discriminative characteristic for each binding site that you identified.<br>3. If the context permits, include any relevant small molecules in the binding site description, location, and target protein.<br>4. Output the list of amino acid residues constituting each of identified binding sites. Use the following notation for defining amino acid residues: <chain_id><res_name><res_id>.<br>4.1 chain_id is an optional chain identifier for each amino acid. Crucially, if the paper specifies a chain ID for a residue (e.g., α, β, γ, A, B, C), you must include that chain ID in the output.<br>4.2 res_name is the amino acid name in either single-letter (A123) or three letter notation (Ala123). Use the same notation as in the paper.<br>4.3 res_id is the integer amino acid number.<br>4.4 Some residues may be followed by a punctuation mark or a space. Any characters that follow them are not part of the residue.<br>Examples:<br>Input Text: "Glu89" -- Extracted Residue: {'chain_id: None, res_name: Glu, res_id: 89'}<br>Input Text: "A134" -- Extracted Residue: {'chain_id: None, res_name: A, res_id: 134'}<br>Input Text: "GLY1204" -- Extracted Residue: {'chain_id: None, res_name: GLY, res_id: 1204'}<br>Input Text: "Ala123 ELC" -- Extracted Residue: {'chain_id: None, res_name: Ala, res_id: 123'}<br>Input Text: "γPhe222" -- Extracted Residue: {'chain_id: γ, res_name: Phe, res_id: 222'}<br>Input Text: "αY45" -- Extracted Residue: {'chain_id: α, res_name: Y, res_id: 45'}<br>Input Text: "Tyr101C" -- Extracted Residue: {'chain_id: C, res_name: Tyr, res_id: 101'}</td>
</tr>
</table>

Table 1. Extraction step prompts before and after optimization.
